# Isolation and Identification of P(3HB)-Degrading Bacteria from Malaysian Environment

**DOI:** 10.1101/2024.10.18.619168

**Authors:** Umber Zahra, A.A Amirul, Ali Nawaz, Hamid Mukhtar

## Abstract

The aim of this study was to identify Poly(3-hydroxybutyrate) [P(3HB)] degrading bacterial strains in Penang Malaysia. Samples were collected from various locations in Penang. Five isolates (SMA-1, SMB-2, SMC-3, SMD-4, and SME-5) with highest P(3HB) degradation indices were isolated through meticulous screening. These gram-negative bacilli were characterized first morphologically and then through scanning electron microscopy. Growth profiles of the isolates revealed that at 24 hrs, SME-5 showed the highest growth rate. Maximum depolymerase enzyme activity for all five isolates was observed after 72 hrs. 16S rDNA sequencing identified all isolates as *Comamonas testosteroni*. This study provides insights into the growth characteristics and extracellular depolymerase activity of P(3HB)-degrading bacteria isolated from the Malaysian environment. It highlights the potential of *Comamonas testosteroni* strains as P(3HB) bioplastic degraders in diverse habitats.

## INTRODUCTION

Poly(3-hydroxybutyrate) [P(3HB)] is a naturally occurring polymer that is fully biodegradable in nature through the action of extracellular depolymerase enzymes secreted by certain microorganisms (Jendrossek & Handrick, 2002). Some microorganisms store P(3HB) intracellularly in the form of granules, in the presence of excess carbon (Kahar *et al*., 2004). However, these granules are released into the environment upon bacterial cell lysis. Extracellular P(3HB) depolymerase enzymes, released by P(3HB) biodegrader microorganisms, convert P(3HB) into water soluble intermediates, which in turn is utilized by these microorganisms as energy source for their growth (Amara & Moawad, 2011).

P(3HB) biodegraders are ubiquitous in the environment i.e. they can be found in activated sludge, compost, soil, as well as in marine and freshwater. P(3HB) degradation activity is highest in anaerobic sewage, whereas, lowest in seawater. The common microbial genera that are known for producing extracellular depolymerase enzymes include *Comamonas sp*., *Pseudomonas sp*., *Alcaligenes sp*. and *Cupriavidus sp*. The fungal groups that are observed for their extracellular depolymerase activities are *Ascomycetes, Basidiomycetes, Deuteromycetes, Mastigiomycetes* and *Myxomycetes* (Muhammadi *et al*., 2015).

Soil holds the capacity to nurture a variety of P(3HB)-degrading bacteria as it is the natural habitat for them (Boyandin et al., 2013). Anderson et al. (1990) noticed that Polyhydroxyalkanoates (PHAs) degraders are quite ubiquitous and the excess availability of P(3HB) polymers triggers them to utilize it as a carbon source.

The identification of P(3HB) biodegrader microorganisms has evolved beyond the traditional techniques. Incorporation of advanced techniques like electron microscopy and 16S rDNA sequencing helps in a more comprehensive analysis. Notably, a recent study conducted by Vigneswari et al. (2015), has elucidated the efficacy of 16S rDNA sequencing in identification of P(3HB)-degrading Bacteria.

The applications of P(3HB) include the development of implanted medical devices particularly for dental surgery, orthopedic, hernioplasty and skin treatments. Moreover, P(3HB) has been used in the development of therapeutic products such as in microspheres and microcapsules for drug delivery. Besides P(3HB) application in development of medical devices and sustained drug delivery, P(3HB) has currently been used to produce enzyme activators or inhibitors for drafting physiological models. P(3HB) presents minimal inflammatory tissue reaction under implantation and can prolong local enzyme activation or inhibition in vivo (Bonartsev*et al*., 2007).

## MATERIALS AND METHODS

### Isolation and Screening of P(3HB)-degrading Bacteria

Eight water samples and two soil samples were collected from various locations in Penang, Malaysia. About 1g of soil sample and 1ml of water sample were added into the 100ml flask containing P(3HB) selective medium and incubated for 24 hours according to the method of Azami et al. (2017). After 24 hours, a serial dilution was performed. A volume of 0.1ml with dilution of 10^-4^, 10^-5^, 10^-6^ were streaked onto the P(3HB) agar plates and incubated at 30°C for 3 days to observe for the halo zone formation. Colonies forming halo zones were selected and purified. Pure colonies were stabbed onto the P(3HB) plates, incubated at 30°C for 3 to 5 days to calculate the degradation index. Degradation indices were calculated by dividing Halo zone size by colony size (Ramachandran & Abdullah, 2010).

### Identification of P(3HB)-degrading bacteria

Bacterial identification was carried out using morphological and molecular analysis for the isolates with the highest degradation index. In morphological identification, the colony characteristics were studied and gram-staining was performed. Scanning Electron Microscopy (SEM) was also carried out to identify the shape of the bacteria. 16S rDNA was extracted for each isolate. The extracted DNA was subjected to PCR amplification. The products obtained after PCR were sent to Integrated DNA Technology for DNA purification and DNA sequencing. For molecular analysis, purified 16S rDNA amplicons were sequenced. The 16S rDNA sequences obtained were subjected to BLAST against reference sequences in the Genbank database (NCBI), to observe homology. The results from BLAST were put into Jalview to construct phylogenetic trees representing closest relatives (Flothet. al., 2000).

### Production of P(3HB) depolymerase enzyme

The isolated P(3HB)-degrading bacteria were incubated at 30°C in a 200 rpm orbital shaker for 120 hours, after overnight enrichment in NR broth. Samples were taken every 24 hrs and centrifuged. The pellet was used to determine bacterial growth by measuring optical density at 540nm (Azami et al., 2017). To access the extracellular depolymerase activity, 100ul of supernatant containing crude depolymerase enzyme was added into Eppendorf tube along with 400ul sodium-glycine buffer and 500ul of 0.08% of P(3HB) substrate. The Eppendorf was incubated at 40°C for 30 minutes and the optical density was taken at 540nm (Gudmalwar and Kamble, 2014). The enzyme activity was determined using the formula below:

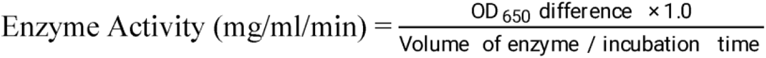

## RESULTS & DISCUSSION

### Isolation and Screening of P(3HB)-degrading bacteria

Bacteria isolated from various samples were screened out by observing the halo zone formation. From the screening, five bacterial isolates (SMA-1, SMB-2, SMC-3, SMD-4, SME-5) showed good formation of the halo zone on the P(3HB) agar. SMB-2 showed the highest degradation index whereas SMC-3 has the lowest degradation index. Degradation index indicates the P(3HB) polymer degrading capability. (Muhammadi *et al*., 2015). Only these five isolates were further studied for enzyme production and growth profile.

### Identification of P(3HB)-degrading bacteria

In colony morphology, four isolates (SMB-2, SMC-3, SMD-4, SME-5) appeared off-white or white in color with rounded shape, while one isolate (SMA-1) appeared to have wrinkled shape. Under compound microscope at 100× magnification with oil emulsion technique, all isolates stained pink in color representing Gram negative behavior. These five isolates were further analyzed under Scanning Electron Microscope at magnification 10,000× for evaluation of their shape. All the isolates were identified rod-shaped as seen in figure 1.

**Figure 1:**
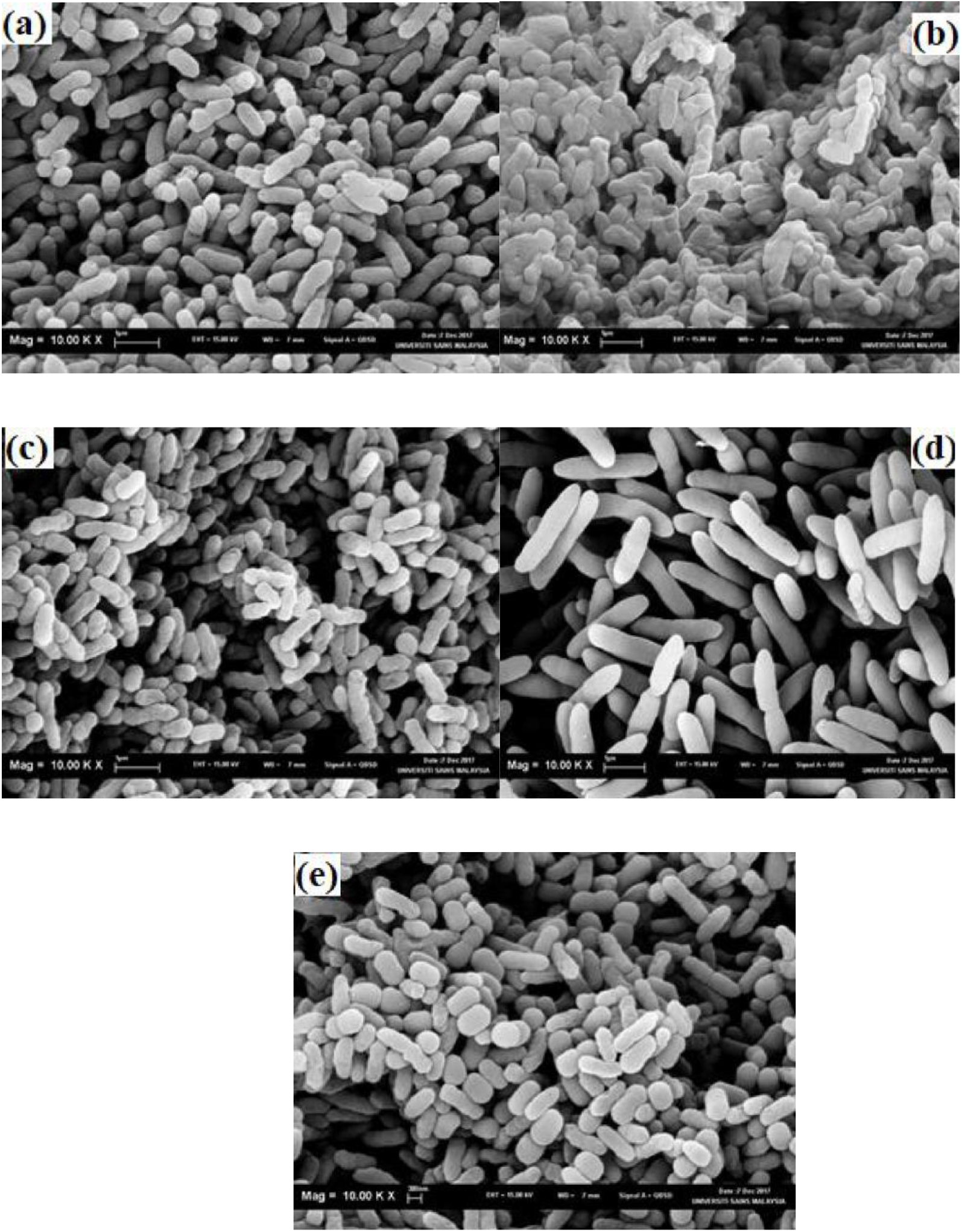
Electron micrographs of five best P(3HB)-degrading Bacterial isolates i.e. (a) SMA-1, (b) SMB-2, (c) SMC-3, (d) SMD-4, (e) SME-5.

### Growth profile and Estimation of enzyme activity

Figure 2 shows the growth profile of the five P(3HB)-degrading bacterial isolates. The maximum growth was observed at the 24 hours of incubation. The growth of the bacteria decreased upon further incubation. Among the isolates, the highest growth (3.228 OD) was shown by SMA-1 and the lowest (1.761 OD) was observed for SMC-3. Bacteria secrete the extracellular hydrolytic enzyme that degrades the P(3HB) polymer into the compounds that are soluble in water and can be utilized by the bacteria as their energy source (Azami*et al*., 2017). Stationary phase of growth was observed after 72 hrs.

**Figure 2:**
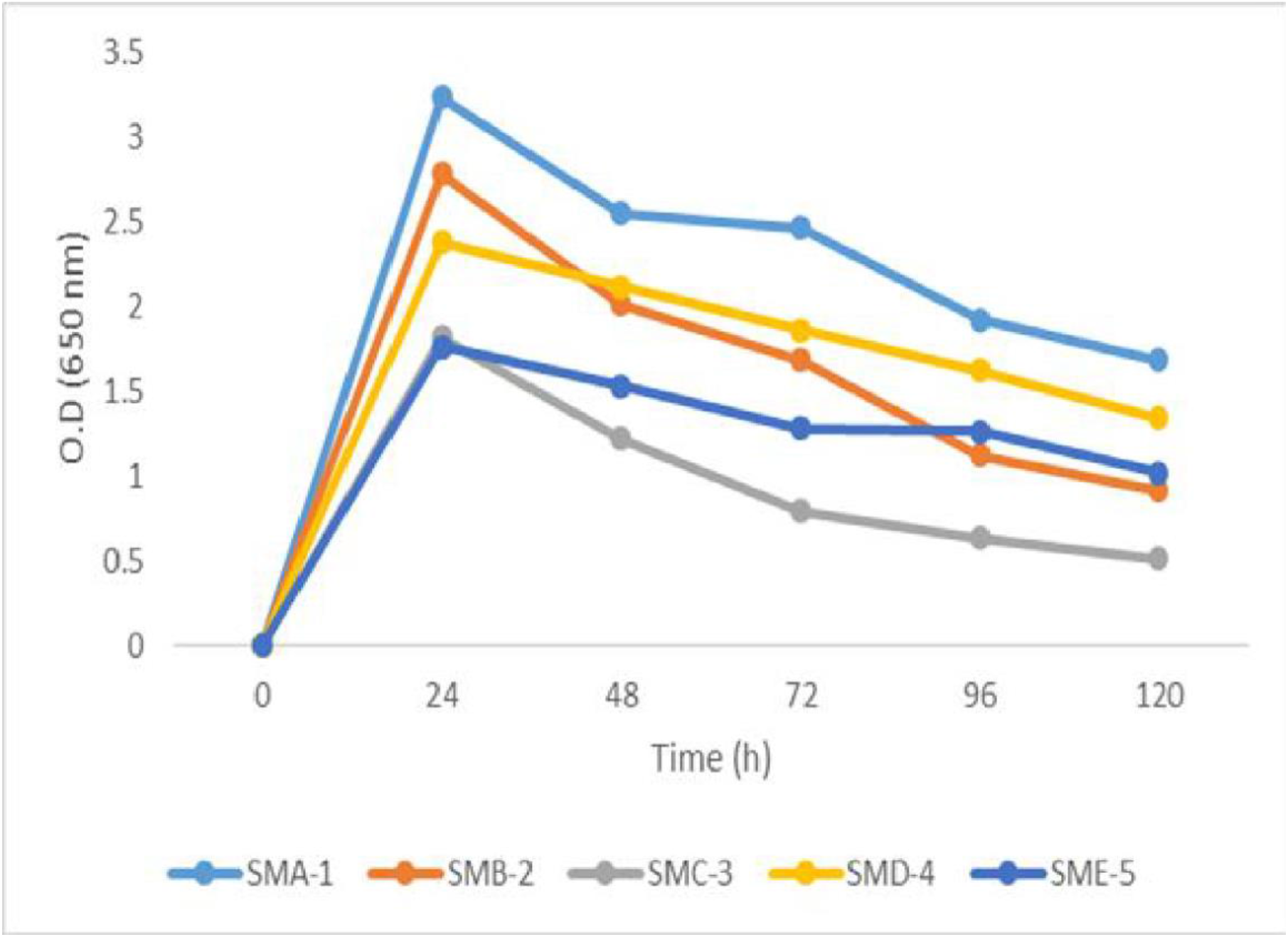
Growth profile of five best P(3HB)-degrading Bacterial isolates.

Figure 3 shows the extracellular depolymerase enzyme activity (Units/ml/min) of all five isolates. All strains exhibited the same pattern where the maximum enzyme activity was achieved at 72 hrs and maintained thereafter. This perfectly aligns with the findings of Vigneswari*et al*. (2015), who also observed the highest enzyme activity at 72 hours in their study. However, it has been previously reported that with the increase in incubation time, the enzyme activity decreases likely due to the inhibition of enzyme depolymerase (Manna and Paul 2000). But we did not see a significant decline in enzyme activity even until the 120 hours mark.

**Figure 3:**
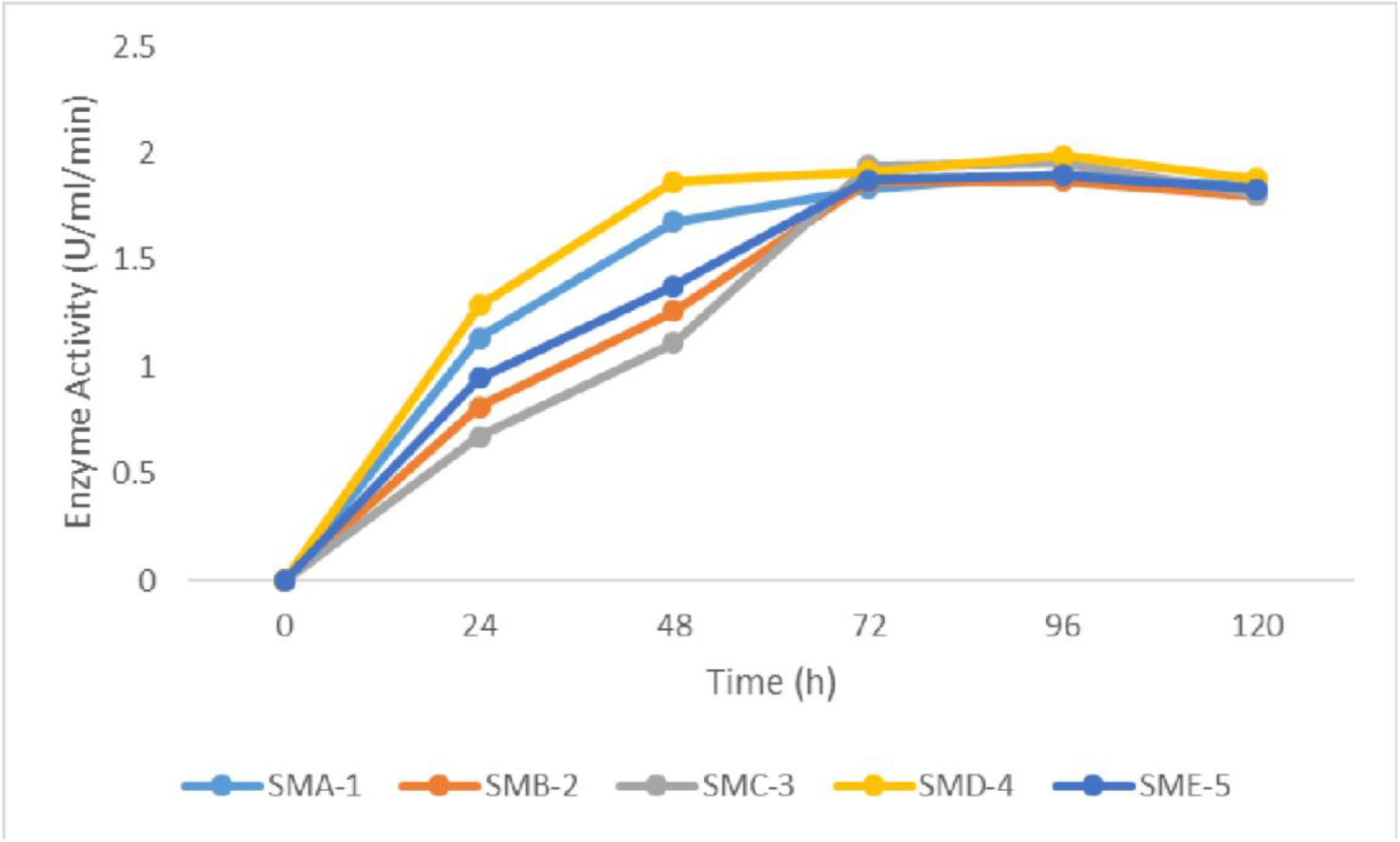
Depolymerase activity of five best P(3HB)-degrading Bacterial isolates.

### Molecular Identification

Results from BLAST indicated that 16S rDNA sequences of all strains exhibit 99% similarity to *Comamonas testosteroni*. This strongly suggests that our isolates – SMA-1, SMB-2, SMC-3, SMD-4 and SMD-5, are strains of *Comamonas testosteroni*, which is a popular bioplastic degrader microorganism.

## CONCLUSION

From this study, it can be concluded that P(3HB) bioplastic-degrading bacteria are ubiquitously found in various locations of Penang, Malaysia. These bacteria mostly belong to a common group, with slight differences in their characteristics and extracellular P(3HB) depolymerase activity. However, it must be noted that this study was limited by a small sample size. In the future, a broader study covering larger sample sizes can validate our results and provide better insights into the characteristics of P(3HB) degrading microorganisms.

